# Collagen organization and structure in *FLBN5*^-/-^, mice using label-free microscopy: implications for pelvic organ prolapse

**DOI:** 10.1101/2024.01.31.578106

**Authors:** Christian M. Jennings, Andrew C. Markel, Mari J.E. Domingo, Kristin S. Miller, Carolyn L. Bayer, Sapun H. Parekh

## Abstract

Pelvic organ prolapse (POP) is a gynecological disorder described by the descent of superior pelvic organs into or out of the vagina as a consequence of disrupted muscles and tissue. A thorough understanding of the etiology of POP is limited by the availability of clinically relevant samples, restricting longitudinal POP studies on soft-tissue biomechanics and structure to POP-induced models such as fibulin-5 knockout (*FBLN5*^*-/-*^) mice. Despite being a principal constituent in the extracellular matrix, little is known about structural perturbations to collagen networks in the *FBLN5*^*-/-*^ mouse cervix. We identify significantly different collagen network populations in normal and prolapsed cervical cross-sections using two label-free, nonlinear microscopy techniques. Collagen in the prolapsed mouse cervix tends to be more isotropic, and displays reduced alignment persistence via 2-D Fourier Transform analysis of images acquired using second harmonic generation microscopy. Furthermore, coherent Raman hyperspectral imaging revealed elevated disorder in the secondary structure of collagen in prolapsed tissues. Our results underscore the need for *in situ* multimodal monitoring of collagen organization to improve POP predictive capabilities.

## 1. Introduction

Pelvic Organ Prolapse (POP) is a multi-etiological disorder characterized by the descent of pelvic organs through the pelvic floor. Approximately 50% of women develop POP in their lifetime [1] POP negatively affects a woman’s quality of life including sexual activity and symptoms of urinary incontinence, bowel incontinence, and pain [2]. Although the primary antecedents to POP are birth-associated injury to the levator ani muscles, age, and body mass index [3], hereditary connective tissue disorders (e.g., Marfan and Ehlers-Danlos syndromes) can cause POP [4] – necessitating studies aimed at elucidating the multifactorial underpinnings of connective tissue dysfunction in POP.

Due to their similar reproductive anatomy to humans [5], rodent models of elastinopathies are used to study POP [6]. One such model is fibulin-5 homozygous knockout mice (KO). In the KO model, over 90% of mice develop severe POP in six months [4]. Fibulin-5 is a glycoprotein produced in smooth muscle cells, fibroblasts, or vascular endothelial cells [7] that is responsible for proper elastic fiber formation [8]. Elastic fiber components such as elastin, fibulins, and fibrillins are critical to proper extracellular matrix (ECM) construction, contributing mechanical compliance and elasticity to the reproductive tract [9]. For example, loss of elastic fibers in arterial walls decreases collagen fiber undulation that affects the collagen fiber’s stiffening characteristics [10]. As a core component of the ECM, collagen fibers undergo greater loads without elastic fiber support. Consequently, collagen can be remodeled in the KO reproductive tract. Clark-Patterson *et al*. demonstrated significantly reduced collagen alignment ratio with respect to the circumferential and axial axes in KO murine vaginas compared to *FBLN5*^*+/-*^ (HET) murine vaginas [9]. Furthermore, Budatha *et al*. reported thin and contorted collagen fibers in murine KO vaginas [11]. In these studies, the bulk tissue and collagen were visualized using histological staining and Picrosirius Red coupled with polarized light microscopy. However, studies investigating collagen orientation and structural perturbations in KO mice cervices are limited.

Like polarized light imaging, second harmonic generation microscopy (SHG) is used to image collagen, albeit in a label-free capacity. SHG is a nonlinear, frequency-doubling optical phenomenon that occurs in non-centrosymmetric materials. In biological tissue, SHG signal arises primarily from type I collagen in supramolecular assemblies [12] – enabling molecularly specific, high-resolution collagen imaging. SHG imaging has demonstrated ability as a modality to study cervical collagen: using surgical biopsies of human cervices, Narice *et al*. observed a significant difference in collagen alignment in pre- and post-menopausal women via SHG imaging [13]. Moreover, Fourier-Transform (FT) SHG analysis was used to observe that collagen orientation varied along the transverse plane but not in the longitudinal plane of wild-type rat cervices [14]. The specificity and resolution of SHG imaging enable quantification of diameter, length, and curvature (amongst others) of individual collagen fibers and bulk collagen network characteristics such as area ratios and alignment. Although outside the scope of this paper, comprehensive reviews of collagen SHG imaging analysis techniques are available [15–17].

The collagen orientation index (COI) is a parameter extracted from FT SHG analysis that permits quantification of the degree of collagen anisotropy within a region of interest (ROI). Briefly, the COI is a ratio of the ellipticity of the 2D power spectrum of a collagen SHG image assessed via ellipse fitting. SHG FT analysis can reveal collagen fibers’ orientation and periodicity [18–20]. 2D FT analysis has been used to quantify collagen orientation in the dermis in response to uniaxial loads [18], in the human optic nerve head [21], and posterior cruciate ligament [22], amongst other tissues.

Coherent Raman hyperspectral imaging is a label-free, diffraction-limited technique capable of capturing the structural motifs of proteins [23–25]. To combat the low efficiency of spontaneous Raman microscopy, broadband coherent anti-Stokes Raman scattering (BCARS) microscopy employs multiple electromagnetic fields to coherently drive and probe a broad bandwidth of vibrational modes at their resonance frequencies within a focal volume [26]. Concerning collagen, coherent Raman imaging can detect proline-rich and -poor regions, hydroxyproline, and triple-helical structure in the amide I and III bands [27]. To the best of the authors’ knowledge, there is no extensive study of collagen structural or molecular changes in the cervix of murine POP models.

In an established mouse model of POP [4, 6, 28], we examined collagen organization and molecular structure in the cervix’s internal and external os (**Fig. 1 A** and **B**) using label-free SHG and BCARS imaging. In axial cross-sections, we imaged endogenous tissue fluorescence via two-photon microscopy and collagen with SHG microscopy (**Fig. 1 C-E**). We observed a significant difference in collagen anisotropy via COI in diseased and heterozygous murine cervices. Using the local mean absolute angular difference (LMAAD), a further quantification of the orientation angle from 2D power spectrum analysis, we observed a significant difference in the spatial persistence of the collagen orientation in the HET and KO internal and external os. Furthermore, collagen in the internal os and external os of KO mice displayed a significant difference in structural disorder compared to HET mice via BCARS micro-spectroscopy.

**Fig. 1.**
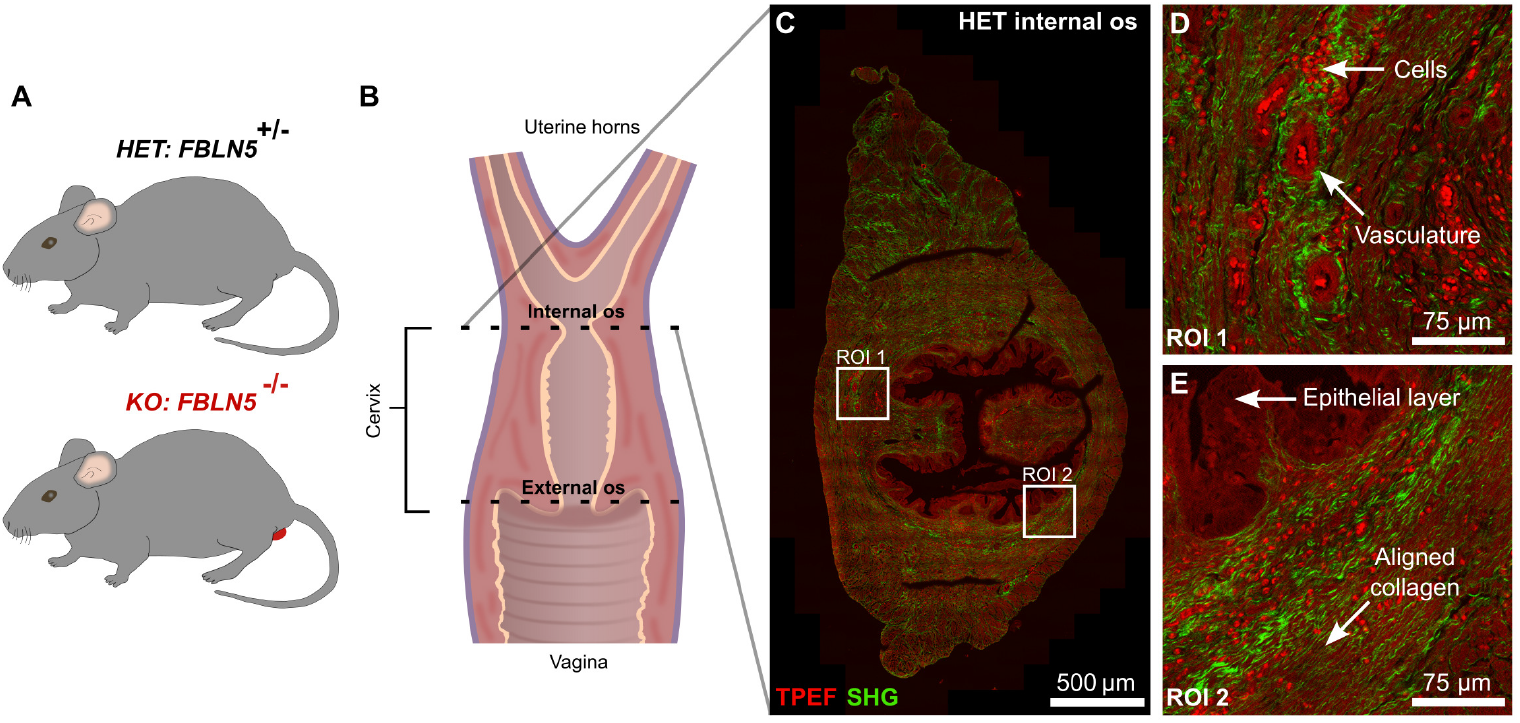
In this study, we compared endocervical collagen organization and structure in heterozygous fibulin-5 (HET) mice and homozygous fibulin-5 knockout (KO) mice with pelvic organ prolapse (**A**). From excised murine reproductive tracts, axial cross-sections 5 µm in thickness were made at the external os and internal os of the cervix (**B**). Each cross-section was imaged using two-photon excited fluorescence (TPEF) of endogenous fluorophores and second harmonic generation (SHG) of collagen. A TPEF and SHG stitched image of a HET internal os cross-section is shown in **C**, highlighting collagen alignment perpendicular to the cervical canal. The image was formed using high-resolution tiles to resolve tissue structures and thin collagen fibers (**D** and **E**).

## 2. Materials and Methods

### 2.1. Animal care

The mouse care and maintenance protocol used was in accordance with Tulane University’s Institute Animal Care and Use Committee. Female mice used in this study were produced from female and male *FBLN5*^*+/-*^ (HET) mice on mixed background (C57BL/6 x 129SvEv). All mice were graded according to the Mouse Pelvic Organ Prolapse Quantification system [29]. *FBLN5*^*-/-*^ (KO) mice with grade 2 or grade 3 perineal bulge were included in the experimental group representing prolapse while grade 0 HET mice were included in the control group. Four mice per group between three and eight months of age were used in this study. All mice were sacrificed via CO_2_ asphyxiation with cervical dislocation as a secondary form of euthanization. Following euthanization, reproductive tract samples were excised from each mouse by making incisions at the uterine bifurcation and the vagina.

### 2.2. Tissue cross-section preparation

Reproductive tract samples were washed with HBSS, fixed in formalin for 24 hours, and embedded in paraffin for histological sectioning. Using a cryotome, five-micron thick tissue cross-sections were cut every 0.5 mm along the reproductive tract and placed on a microscopy slide. The external os of the cervix was identified in slides proximal to the vagina by the presence of the vaginal fornices lateral to the cervical os, while the internal os of the cervix was identified in slides distal to the uterine horns by the presence of only the cervical os (no fornices). Before imaging, the samples were deparaffinized and rehydrated. The samples were washed twice in xylene, once in a xylene and 100% ethanol solution (1:1 v/v), twice in 100% ethanol, once in 95% ethanol, once in 70% ethanol, once in 50% ethanol, and 1X phosphate buffered saline. The duration of each wash step was three minutes. A cover glass (22 x 22 mm, #2; VWR, Radnor, PA, USA) offset with double-sided tape spacer, constituted a chamber sealed using Valap (1:1:1 mixture of petroleum jelly, lanolin, and paraffin) to reduce sample dehydration. Samples were kept hydrated at 4°C between imaging sessions.

### 2.3. TPEF and SHG imaging

We examined collagen morphology and alignment using endogenous two-photon excited fluorescence (TPEF) and second harmonic generation (SHG) from collagen. Images were acquired using a laser-scanning, confocal microscope (FV3000, Olympus, Tokyo, Japan). Multi-photon excitation was generated using an ultra-fast laser (InSight X3, Spectra-Physics, Milpitas, CA, USA) tuned to 800 nm and focused to the sample plane with a water-immersion, 60X, 1.1 NA objective (LUMFLN, Olympus, Tokyo, Japan). The excitation beams were linearly polarized as circularly polarized light reduced the overall SHG signal and made minimal impact on the observed collagen structure (SI Fig. 1). Image tiles (512 x 512 pixels; 0.276 µm/pixel) were averaged twice per line, and acquired using a 1.5X optical zoom at 12 bits/pixel. Tiles were stitched into a whole cross-section image using an onboard Olympus Correcting Algorithm. SHG and TPEF emission were acquired in the epi-configuration. The signals were isolated from the 800 nm excitation beam with a 680 nm short-pass filter (FF01-680/SP-25, Semrock, West Henrietta, NY, USA). A dichroic 425 nm long-pass filter (DLMP425R, Thorlabs, Newton, NJ, USA) splits the TPEF and SHG signal, and the SHG signal is filtered further by a 405 ± 5 nm band-pass filter (FBH405-10, Thorlabs, Newton, NJ, USA), before detection by home-built photomultiplier tubes (PMTs). The PMTs’ gains were kept constant across each sample.

### 2.4. COI and LMAAD post-processing

We quantified the morphology and spatial persistence of collagen fiber alignment through the collagen orientation index (COI) and local mean absolute angular difference (LMAAD) metrics, respectively. Using the stitched SHG image, we converted the image to 8 bits/pixel and applied a Gaussian blurring filter (*σ*_r_ = 0.25 µm) in ImageJ (2.14, National Institute of Health, USA). From this smoothed SHG image, we produced a binary collagen mask using ImageJ’s mean threshold. The smoothed SHG image and the mask were inputs to a custom MATLAB (2023a, MathWorks, Natick, MA, USA) script to perform COI and LMAAD processing.

The COI and LMAAD metrics rely on ellipse fitting of a binarized power spectrum (**Fig. 2 C** and **D**). Power spectra were calculated from the smoothed SHG image in 64 x 64-pixel windows in locations where the SHG mask had values of one in 70% of its area. Each power spectra’s zero-frequency component was centered and a log_10_ transform was applied. Each power spectrum was binarized using half-the-max intensity plus 0.2 of the power spectrum’s radial average. We fit the binarized power spectrum with an ellipse to ascertain the ellipse’s major and minor axes’ length and its orientation angle in degrees. Each window’s COI was quantified as [1 – (minor axis length / major axis length)]. Using the orientation angle (between -90° and 90°) of each ellipse’s major axis, we calculated the natural logarithm of the mean absolute angular difference between the center window (“0”) and the adjacent windows (“1”), the windows two windows away from the center (“2”), and the windows three windows away from the center (“3”) (**Fig. 3 A-D**). We subtracted 180° from absolute angular differences exceeding 90° to prevent false large angular differences. Windows without a power spectrum by virtue of the collagen mask exclusion were omitted from the LMAAD calculation.

**Fig. 2.**
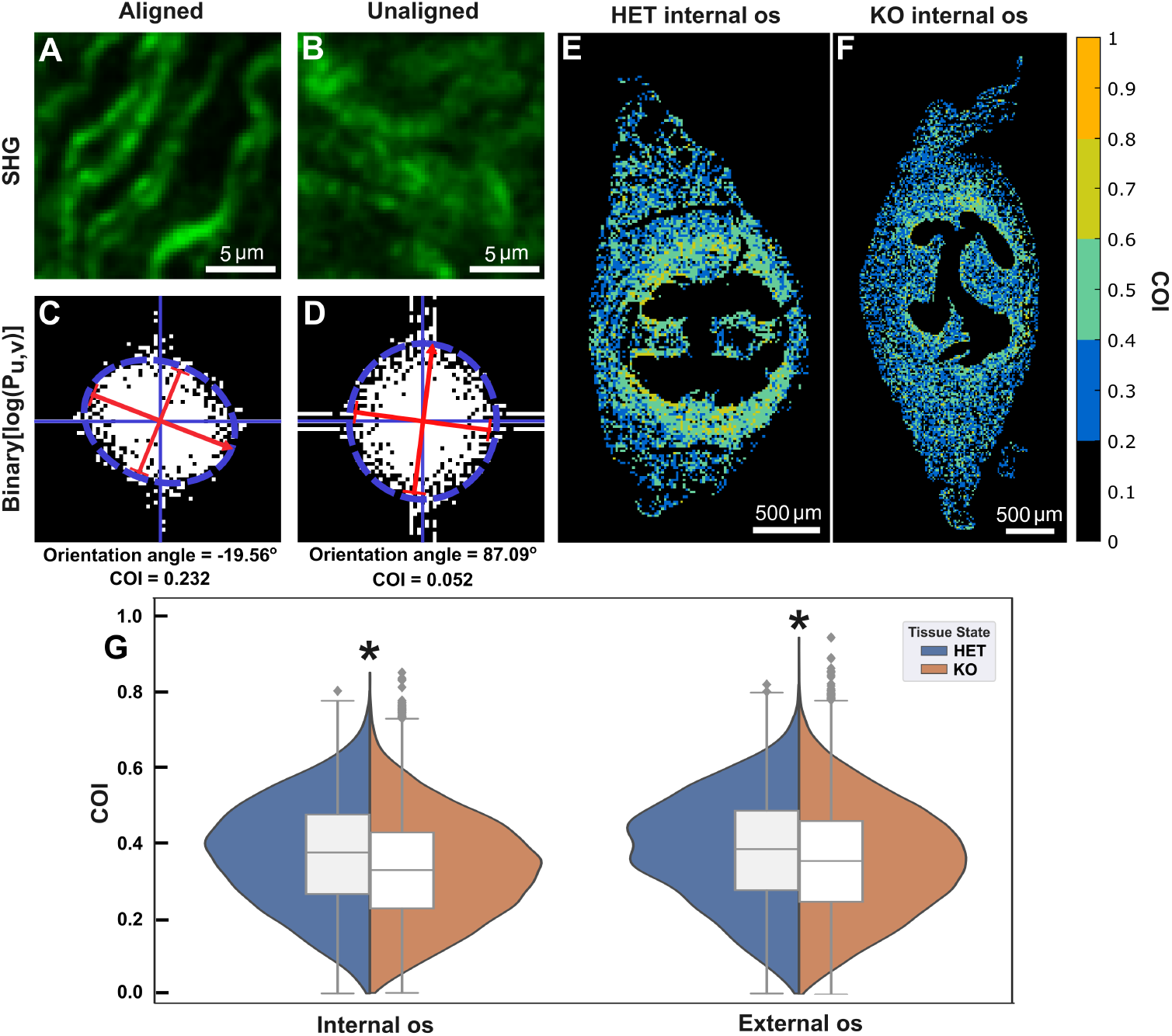
The collagen orientation index (COI) indicates the degree of collagen anisotropy within a selected window. The binarized power spectrums of anisotropic (**A**) and isotropic (**B**) collagen windows reveal distinct spatial frequency distributions (**C** and **D**). By fitting an ellipse to the power spectral densities, we extract the collagen orientation index as one minus the ratio of the minor axis length to the major axis length. More aligned collagen (**C**) has a greater COI than isotropic collagen (**D**). Furthermore, the orientation angle of the major axis (-90° and 90°) is extracted from ellipse fitting for LMAAD quantification. The COI was computed on distinct 64 x 64-pixel windows across every SHG image. The spatial COI values for the HET internal os are displayed in **E**, highlighting greater COIs for the relatively more anisotropic collagen near the cervical canal. (**F**) The HET internal (N=4, n=13,716) and external os (N=4, n=17,430) display significantly greater mean COIs compared to the KO internal (N=4, n=31,272) and external os (N=4, n=51,727). ^*^ Indicates a P < 0.001 via two-tailed Mann-Whitney-Wilcoxon test.

**Fig. 3.**
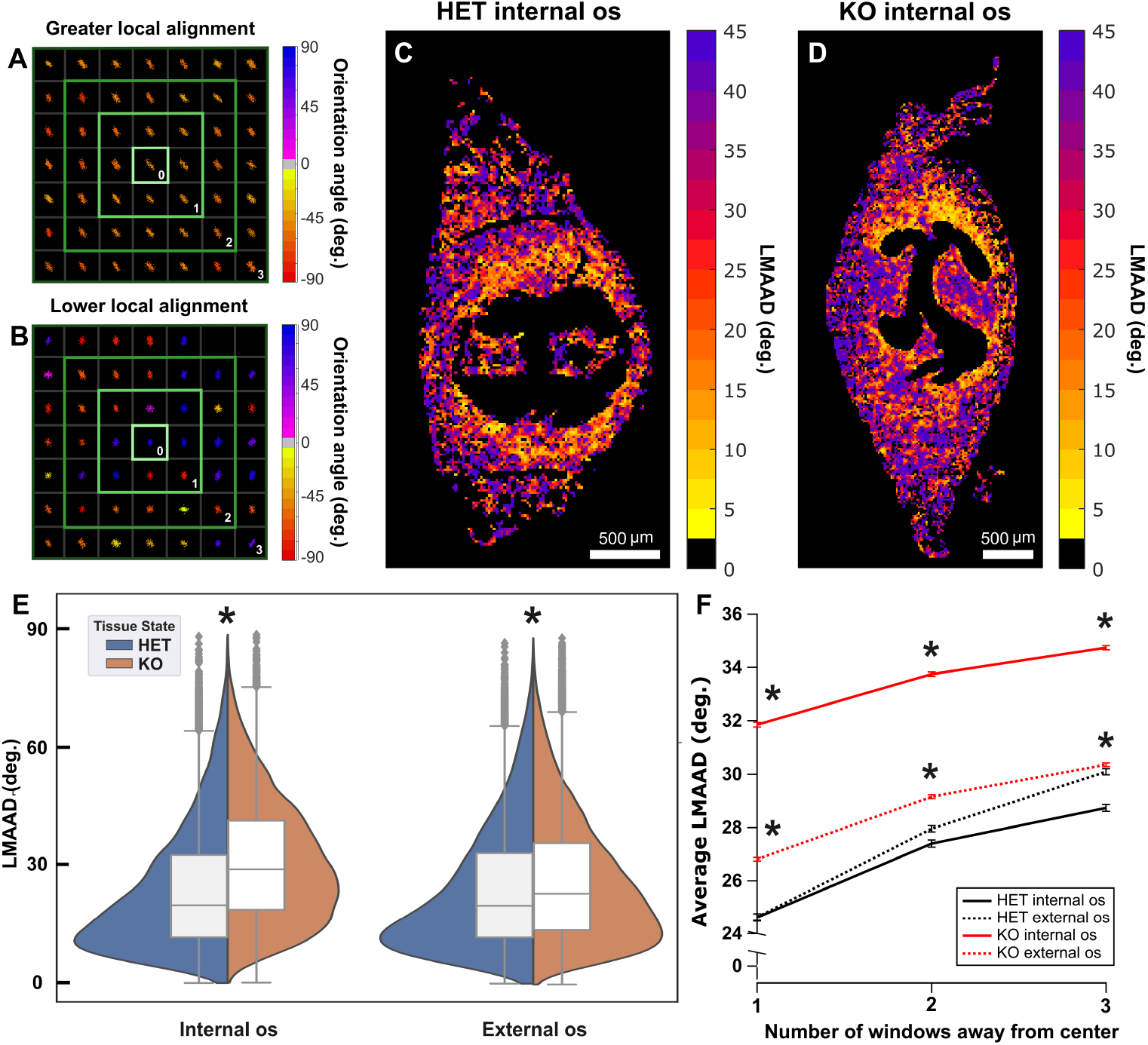
The local mean absolute angular difference (LMAAD) quantifies the persistence of collagen orientation. Using each window’s (64 x 64 pixels) major axis orientation angle, we calculate the mean absolute angular difference between the center window (“0”) and the adjacent windows (“1”), the windows one window from the center (“2”), and windows two windows from the center (“3”) (**A** and **B**). The result is assigned to the center window. (**D** and **C**) Applying this processing (“1”) to one of the HET and KO internal os samples indicates low LMAAD near the cervical canal. Violin plots of compiled LMAAD distributions for the adjacent windows (“1”) are highlighted in E (HET: int. os, n=12,913; ext. os, n=15,815; KO: int. os, n=29,989; ext. os, n=48,970). (**F**) The HET internal (N=4) and external os (N=4) display significantly different LMAAD compared to the KO internal (N=4) and external os (N=4) as a function of window distance. ^*^ Indicates a P < 0.001 via two-tailed Mann-Whitney-Wilcoxon test.

### 2.5. BCARS hyperspectral imaging

We used a home-built broadband coherent anti-Stokes Raman scattering microscope to probe collagen structural motifs. For excitation, a Nd:YAG microchip laser generates nanosecond pulses with a repetition rate of approximately 1 MHz at 1064 nm and a broadband supercontinuum that ranges from 1100 – 2400 nm (Opera HP, Leukos, Limoges, France). The beams were coupled in the sample plane via a 100X, 0.85 NA objective (LCPLN100XIR, Olympus, Tokyo, Japan). In a transmission configuration, the signal was collected with a 20X, 0.4 NA objective (M-20X, MKS Newport, Andover, MA, USA). Signal was measured with a spectrometer (IsoPlane 160, Teledyne Princeton Instruments, Trenton, NJ, USA) and a back-illuminated, deep-depletion CCD (Blaze 1340 x 400 HS, Teledyne Princeton Instruments, Trenton, NJ, USA). We used a 1600 nm short-pass filter (84-656, Edmund Optics, Barrington, NJ, USA) in the supercontinuum path to limit thermally produced bubbles during acquisition. Hyperspectral images were acquired by stage-scanning the tissue cross-sections in 50 x 50-pixel tiles at a step size of 0.40 µm/pixel and an integration time of 40 ms/pixel. Quantitative comparisons of collagen structure operated on a stitched hyperspectral image (8 x 8 tiles, 160 µm^2^). At a maximum, stitched images were composed of 15 x 15 tiles (300 µm^2^ hyperspectral image) (**Fig. 4 B**). Using the merged TPEF and SHG images as a guide, CARS imaging for all samples was relegated to collagen at or near the basement membrane.

**Fig. 4.**
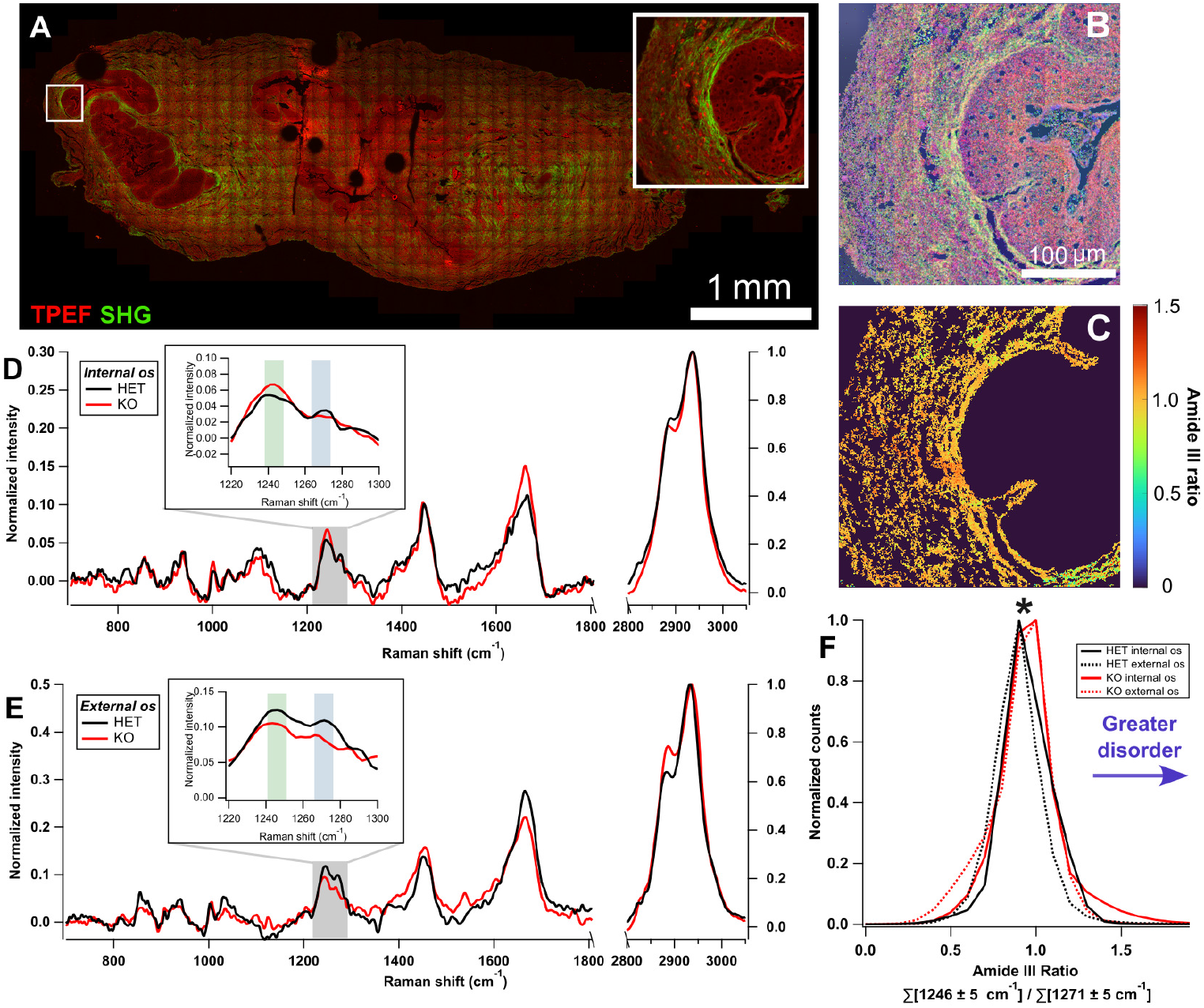
The structural order of collagen in internal os (HET, N=4; KO, N=4) and external os (HET, N=3; KO, N=4) cross-sections probed using BCARS microscopy. (**A**) TPEF and SHG images are used to locate subepithelial collagen for BCARS imaging. (**B**) Molecular pseudocolor of the ROI of the whole image in **A**. Red is protein signal arising in the amide I (C=O stretch at 1659 cm^-1^), green is assigned to the amide III of collagen (1246 cm-1), and blue is assigned to phenylalanine (1002.2 cm^-1^). (**C**) Using the intensity of amide III of collagen (1246 cm-1) as a mask, the ratio of disordered to ordered collagen is quantified as the sum of 1246 ± 5 cm^-1^ over the sum of 1271 ± 5 cm^-1^. Representative normalized, averaged (5 x 5 pixels) collagen spectra in the internal os (**D**) and external os (**E**). The combined amide III ratio distributions (20 bins, bin size is 0.1) indicate significantly different collagen structural disorder in HET (int. os, n=154,833; ext. os, n=75,365) and KO (int. os, n=128,599; ext. os, n=135,142) tissues (**F**). ^*^ Indicates a P < 0.001 via two-tailed Mann-Whitney-Wilcoxon test.

### 2.6. BCARS post-processing

Using a modified Kramers-Kronig transform for phase retrieval, raw BCARS spectra were transformed into Raman-like spectra for quantitative analysis as reported in previous studies [23, 30]. Then, a second-order Savisky-Golay with a 151-point (approximately Δ396 cm^-1^) smoothing window was applied to the phase retrieved spectra to produce the Raman-like spectra. These operations were performed in Igor Pro (8.04, WaveMetrics, Portland, OR, USA). The Raman-like hyperspectral datasets were exported to MATLAB (2023a, MathWorks, Natick, MA, USA) for all subsequent processing.

Center wavelength dependencies of the spectrometer calibration required a spectral shift of less than or equal to 10 cm^-1^. Consequently, each spectrum was shifted to have its maximum located at 2934.6 cm^-1^, which arises from CH_3_ vibrations [31]. Then, the spectrum in each spatial pixel was computed as a 3 x 3-pixel average to reduce spectral noise, which amounts to a smoothing over 1.44 µm^2^. We then normalized each spectrum by its respective maximum at 2934.6 cm^-1^. We then zeroed the spectrum of all pixels with mean values less than half of each tiled scan’s integrated amide I (1580 – 1700 cm^-1^) intensity; these pixels did not have sufficient material to warrant further analysis. A collagen mask was made by projecting the I_1246_/I_2934_ intensity, followed by a 2D Gaussian blurring filter (*σ* = 0.5), and image binarization via Otsu’s threshold. We then manually removed any remaining scan acquisition artifacts and hot pixels outside the tissue boundary using the merged TPEF and SHG images as a guide. Using the resultant mask, we quantified the degree of collagen disorder through the amide III ratio of disordered collagen intensity (1246 ± 5 cm^-1^) over ordered collagen intensity (1271 ± 5 cm^-1^) at each hot pixel in the image [32]. We removed amide III ratios equaling exactly one as typically these arose from pixels with quite low amide III signal-to-noise ratios.

### 2.7. Statistics

Statistical analysis was performed in RStudio (4.3.1, The R Foundation for Statistical Computing). First, the normality of each distribution was assessed using the Shapiro-Wilk test, and all distributions were found to significantly deviate from a normal distribution (P > 0.05). Therefore, the nonparametric two-tailed Mann-Whitney-Wilcoxon test was used to determine whether HET and KO COI, LMAAD, and amide III ratio distributions were differed significantly (P < 0.001).

## 3. Results

### 3.1. Disrupted collagen alignment in fibulin-5 KO mice across multiple length scales

We quantified the distribution of collagen anisotropy from SHG images of the internal and external os of HET and KO mice cervices. The COI indicates the degree of collagen anisotropy within a selected window by ratiometric quantification of the window’s spatial frequency (binarized 2-D power spectrum) distribution. Regions with anisotropic collagen aligned along a particular direction produce high-frequency components orthogonal to the alignment axis in the 2-D power spectrum (**Fig. 2 A** and **C**). Conversely, isotropic collagen produces a nearly radially equivalent or circular distribution of frequency components in the 2-D power spectrum (**Fig. 2 B** and **D**). The COI of collagen in the basement membrane and near the cervical canal was relatively higher in HET and KO mice than collagen closer to the cross-section periphery (**Fig. 2 E**). Analyzing the COI over the entire tissues, we found that HET and KO COI distributions significantly differed in the internal and external os. HET cervix tissues showed more alignment than those in the KO mice.

Using LMAAD processing, we quantified the persistence of collagen orientation angle as a function of relative window distance. For each fit ellipse, we extracted the orientation angle of the major axis (**Fig. 2 C** and **D**). Locations with greater local collagen fiber alignment have lower LMAAD due to more persistent fiber orientation or congruence over space. Furthermore, plotting an LMAAD map highlights spatial variations in persistent collagen alignment, which showed a lower LMAAD for collagen windows closer to the cervical canal (**Fig. 3 F**). We compared the LMAAD of the HET and KO internal os and external os within adjacent (“1”) windows and one (“2”) and two (“3”) windows away from the center window (**Fig. 3 A** and **B**). Across all window distances, we observe significantly different LMAAD distributions between HET and KO cross-sections (**Fig. 3 E**), with HET mice exhibiting relatively lower mean LMAAD than KO mice (**Table 1**).

**Table 1.**
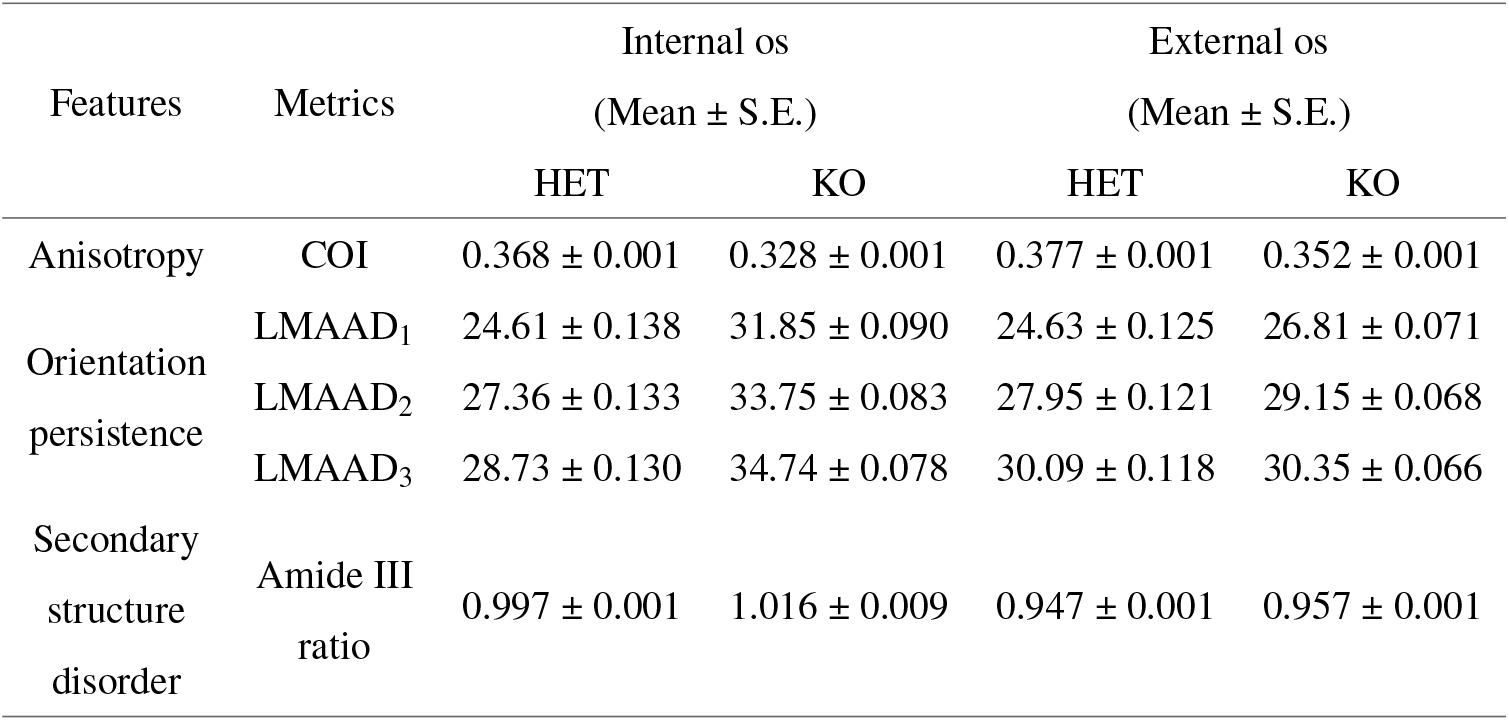
Compiled metrics of collagen morphology and structure in investigated cervical tissues.

| Features | Metrics | Internal os |  | External os |  |
| --- | --- | --- | --- | --- | --- |
| | | (Mean ± S.E.) | | (Mean ± S.E.) | |
|  |  | HET | KO | HET | KO |
| Anisotropy | COI | 0.368 ± 0.001 | 0.328 ± 0.001 | 0.377 ± 0.001 | 0.352 ± 0.001 |
| Orientation persistence | LMAAD<sub>1</sub> | 24.61 ± 0.138 | 31.85 ± 0.090 | 24.63 ± 0.125 | 26.81 ± 0.071 |
| | LMAAD<sub>2</sub> | 27.36 ± 0.133 | 33.75 ± 0.083 | 27.95 ± 0.121 | 29.15 ± 0.068 |
| | LMAAD<sub>3</sub> | 28.73 ± 0.130 | 34.74 ± 0.078 | 30.09 ± 0.118 | 30.35 ± 0.066 |
| Secondary structure disorder | Amide III ratio | 0.997 ± 0.001 | 1.016 ± 0.009 | 0.947 ± 0.001 | 0.957 ± 0.001 |

Moreover, the LMAAD increases with larger distances between windows at both cervical locations in HET and KO mice. This trend is attributed to a decrease in the correlation of fiber orientation as the distance increases between any two collagen windows, as would be expected. However, the rate of LMAAD increase with window distance is approximately equal in HET and KO collagen. The COI and LMAAD analyses reveal a decrease in collagen anisotropy and orientation persistence in KO mice. The HET collagen LMAAD versus distance is consistently lower than for the KO for both internal and external os.

### 3.2. Collagen structural motifs are perturbed in fibulin-5 KO mice

Following observation of perturbed collagen morphology in KO mice, we investigated molecular alterations of KO collagen in the basement membrane (**Fig. 4 A** and **B**) using BCARS hyperspectral imaging. Native, individual collagen fibers are comprised of three polyproline-II helices organized into a triple-helix [33], which is critical for collagen’s biophysical characteristics, particularly its mechanical stiffness [34]. Conversely, disordered collagen resides in a random coil conformation. Representative collagen spectra in HET and KO mice are shown in Figure 4. Several collagen vibrational modes of HET collagen are preserved in the KO collagen. The vibrational modes at 856 cm^-1^ and 937 cm^-1^ arise from proline and hydroxyproline amino acid side chains of collagen as well as C-C vibrations in the collagen backbone [35]. A sharp phenylalanine response is found at 1002.2 cm^-1^. The band at 1457 cm^-1^ arises from CH_2_/CH_3_ deformation and CH_2_ wagging. Furthermore, the amide I (1580 – 1700 cm^-1^) and amide III (1220 – 1300 cm^-1^) bands can be used as vibrational read-outs of collagen secondary structure [27, 36, 37].

We observed a significant difference in the amide III band, which is attributed to collagen’s triple-helical structure [31]. We limited our analysis of collagen secondary structure to the amide III due to the relatively higher collagen specificity of the amide III compared to the amide I (SI Fig. 2) within a focal volume. Vibrational modes at lower Raman shifts of the amide III are attributed to random coil protein secondary structure, whereas *α*-helical protein secondary structures arise in higher Raman shifts of the amide III [32, 38]. Using a ratio of the random coil (1246 ± 5 cm^-1^) and triple-helical (1271 ± 5 cm^-1^) contributions to the amide III band, it is possible to determine relative collagen disorder [39] as we demonstrate using *in vitro* gelatin and collagen I fibers in SI Figure 3. Gelatin, being the denatured form of collagen I, shows a larger ratio of I_1246_/I_1271_ as would be expected since the triple helical structure in gelatin is essentially eliminated. In the cervical cross-sections, HET and KO collagen amide III ratio distributions are significantly different with KO collagen trending towards greater disorder (**Fig. 4 F**).

## 4. Discussion

In this study, we investigated collagen morphology and structure within the cervix of KO and HET mice using label-free microscopic techniques. With statistical significance, we found that KO and HET collagen fibers are from distinct populations regarding their morphology and secondary structure. The collagen in the KO internal and external os tended to be more isotropic, less correlated in orientation angle with respect to neighboring fibers, and more molecularly disordered than HET counterparts. Our observations imply cervical collagen structure and organization can be used to monitor POP development, despite the limited role the cervix has in POP.

The cause of disrupted collagen morphology and structure in the KO mouse cervix is likely multifactorial. KO mice functionally lack elastic fibers that provide mechanical support to soft tissues, limiting the ability of the ECM to recover from deformation in response to applied forces. Without functional and fully assembled elastic fibers, fibrillar collagen, which largely provides the ECM tensile strength, is left to bear increased loading distinct in KO compared to HET soft tissue [9]. Indeed, in vivo and in vitro fibulin-5 HET mouse cervices display significantly greater circumferential contractility than fibulin-5 KO mouse cervices [40]. In addition to mediating elastogenesis, fibulin-5 contains a conserved arginine-glycine-aspartic acid domain that inhibits fibronectin-mediated matrix metalloproteinase-9 (MMP-9) expression [11]. Without fibulin-5, MMP-9 is upregulated therefore impacting collagen fiber degradation and reorganization [41, 42]. Although indirectly, fibulin-5 impacts the mechano-chemical homeostasis of collagen fibers, which potentially establishes a relationship between cervical collagen organization, alignment, and structure and elastic fibers observed in this study. However, most POP studies using fibulin-5 mouse models focus on the vagina and have not included measurements of elastic fibers, collagen, and MMP-9 regulation in anatomy more superior in the reproductive tract.

The COI, LMAAD, and Raman-based collagen quantification can also be applied to cervical collagen in the context of pregnancy. Before delivery, the cervix softens, ripens, and dilates to ensure normal parturition [43]. The consequence of improper timing of cervical ECM remodeling is premature birth [43] that can lead to neonatal morbidity and mortality [44]. Using SHG-based imaging, Atkins *et al*. observed changes in cervical collagen abundance, fiber size, and matrix porosity during gestation that indicated collagen degradation in a preterm mouse model [45]. The COI and LMAAD quantification of bulk cervical collagen organization and orientation persistence may assist identification of irregular cervical ECM and indicate a higher potential for preterm birth. In addition to the morphology of collagen, the degree and type of collagen crosslinking is critical to the cervical mechanical compliance necessary at each stage of gestation [46]. Several studies identified the degree of collagen crosslinking using Raman spectroscopy [47, 48]. Through quantification of cervical collagen crosslinking and structural disorder, Raman-based measurements could be used as hallmarks of incorrect mechanical compliance during gestation.

## 5. Conclusion

In summary, different imaging modalities are needed to observe microstructural and bulk ECM modifications in POP models more holistically. These measurements may clarify the origins of POP and improve clinical outcomes. We demonstrate microstructural changes in prolapsed tissue using nonlinear microscopy that may be preserved in clinically relevant POP presentations. Moreover, we quantified microstructural changes via established (COI) and new imaging processing techniques (LMAAD) that could be clinically useful with e.g. in clinic multiphoton microscopy probes [49–52]. However, improvements to optical fiber-based Raman hyperspectral imaging, specifically amide III signal-to-noise, and TPEF/SHG imaging are necessary to correlate observations in the KO model to patients *in situ*.

## Acknowledgments

C.B. and S.H.P. are grateful to the Research Corporation for Science Advancement Scialog Fellows program for introducing them and initiating this collaboration.

## Data Availability Statement

All data in this paper is available upon reasonable request from the corresponding authors

## Supplemental document

See Supplement 1 for supporting content.

## References

1. M. D. Barber and C. Maher, “Epidemiology and outcome assessment of pelvic organ prolapse,” Int. Urogynecology J. 24, 1783–1790 (2013).

2. L. Carroll, C. O. Sullivan, C. Doody, et al., “Pelvic organ prolapse: The lived experience,” PLOS ONE 17, e0276788 (2022). Publisher: Public Library of Science.

3. T. F. M. Vergeldt, M. Weemhoff, J. IntHout, and K. B. Kluivers, “Risk factors for pelvic organ prolapse and its recurrence: a systematic review,” Int. Urogynecology J. 26, 1559–1573 (2015).

4. P. G. Drewes, H. Yanagisawa, B. Starcher, et al., “Pelvic organ prolapse in fibulin-5 knockout mice: pregnancy-induced changes in elastic fiber homeostasis in mouse vagina,” The Am. J. Pathol. 170, 578–589 (2007).

5. P. A. Moalli, N. S. Howden, J. L. Lowder, et al., “A rat model to study the structural properties of the vagina and its supportive tissues,” Am. J. Obstet. Gynecol. 192, 80–88 (2005).

6. K. Allen-Brady, M. A. T. Bortolini, and M. S. Damaser, “Mouse Knockout Models for Pelvic Organ Prolapse: a Systematic Review,” Int. urogynecology journal 33, 1765 (2022). Publisher: NIH Public Access.

7. D. B. Manders, H. A. Kishore, A. F. Gazdar, et al., “Dysregulation of fibulin-5 and matrix metalloproteases in epithelial ovarian cancer,” Oncotarget 9, 14251–14267 (2018).

8. G. M. Northington, “Fibulin-5: two for the price of one maintaining pelvic support,” The J. Clin. Investig. 121, 1688–1691 (2011). Publisher: American Society for Clinical Investigation.

9. G. L. Clark-Patterson, S. Roy, L. Desrosiers, et al., “Role of fibulin-5 insufficiency and prolapse progression on murine vaginal biomechanical function,” Sci. Reports 11, 20956 (2021). Number: 1 Publisher: Nature Publishing Group.

10. J. Ferruzzi, M. J. Collins, A. T. Yeh, and J. D. Humphrey, “Mechanical assessment of elastin integrity in fibrillin-1-deficient carotid arteries: implications for Marfan syndrome,” Cardiovasc. Res. 92, 287–295 (2011).

11. M. Budatha, S. Roshanravan, Q. Zheng, et al., “Extracellular matrix proteases contribute to progression of pelvic organ prolapse in mice and humans,” The J. Clin. Investig. 121, 2048–2059 (2011).

12. X. Chen, O. Nadiarynkh, S. Plotnikov, and P. J. Campagnola, “Second harmonic generation microscopy for quantitative analysis of collagen fibrillar structure,” Nat. Protoc. 7, 654–669 (2012). Number: 4 Publisher: Nature Publishing Group.

13. B. F. Narice, N. H. Green, S. MacNeil, and D. Anumba, “Second Harmonic Generation microscopy reveals collagen fibres are more organised in the cervix of postmenopausal women,” Reproductive Biol. Endocrinol. : RB&E 14, 70 (2016).

14. T. Lau, H. Sangha, E. Chien, et al., “Application of Fourier transform-second-harmonic generation imaging to the rat cervix,” J. Microsc. 251, 77–83 (2013). _eprint: https://onlinelibrary.wiley.com/doi/pdf/10.1111/jmi.12046.

15. Z. Nejim, L. Navarro, C. Morin, and P. Badel, “Quantitative analysis of second harmonic generated images of collagen fibers: a review,” Res. on Biomed. Eng. 39, 273–295 (2023).

16. Y. Liu, A. Keikhosravi, G. S. Mehta, et al., “Methods for Quantifying Fibrillar Collagen Alignment,” in Fibrosis: Methods and Protocols, L. Rittié, ed. (Springer, New York, NY, 2017), Methods in Molecular Biology, pp. 429–451.

17. Y. Liu, A. Keikhosravi, C. A. Pehlke, et al., “Fibrillar Collagen Quantification With Curvelet Transform Based Computational Methods,” Front. Bioeng. Biotechnol. 8 (2020).

18. S. Nesbitt, W. Scott, J. Macione, and S. Kotha, “Collagen Fibrils in Skin Orient in the Direction of Applied Uniaxial Load in Proportion to Stress while Exhibiting Differential Strains around Hair Follicles,” Materials 8, 1841–1857 (2015).

19. S. Wu, H. Li, H. Yang, et al., “Quantitative analysis on collagen morphology in aging skin based on multiphoton microscopy,” J. Biomed. Opt. 16, 040502 (2011). Publisher: SPIE.

20. X. Zhu, S. Zhuo, L. Zheng, et al., “Quantified characterization of human cutaneous normal scar using multiphoton microscopy,” J. Biophotonics 3, 108–116 (2010). at_eprint: https://onlinelibrary.wiley.com/doi/pdf/10.1002/jbio.200910058.

21. J. K. Pijanka, P. P. Markov, D. Midgett, et al., “Quantification of collagen fiber structure using second harmonic generation imaging and two-dimensional discrete Fourier transform analysis: Application to the human optic nerve head,” J. Biophotonics 12, e201800376 (2019).

22. W. Lee, M. H. Rahman, M. E. Kersh, and K. C. T. Jr, “Application of quantitative second-harmonic generation microscopy to posterior cruciate ligament for crimp analysis studies,” J. Biomed. Opt. 22, 046009 (2017). Publisher: SPIE.

23. S. H. Parekh, Y. J. Lee, K. A. Aamer, and M. T. Cicerone, “Label-Free Cellular Imaging by Broadband Coherent Anti-Stokes Raman Scattering Microscopy,” Biophys. J. 99, 2695–2704 (2010).

24. F. Fleissner, M. Bonn, and S. H. Parekh, “Microscale spatial heterogeneity of protein structural transitions in fibrin matrices,” Sci. Adv. 2, e1501778 (2016).

25. S. Kumar, Y. Wang, M. Hedayati, et al., “Structural control of fibrin bioactivity by mechanical deformation,” Proc. National Acad. Sci. 119, e2117675119 (2022). Company: National Academy of Sciences Distributor: National Academy of Sciences Institution: National Academy of Sciences Label: National Academy of Sciences Publisher: Proceedings of the National Academy of Sciences.

26. C. H. Camp, Y. J. Lee, J. M. Heddleston, et al., “High-Speed Coherent Raman Fingerprint Imaging of Biological Tissues,” Nat. photonics 8, 627–634 (2014).

27. M. G. Martinez, A. J. Bullock, S. MacNeil, and I. U. Rehman, “Characterisation of structural changes in collagen with Raman spectroscopy,” Appl. Spectrosc. Rev. 54, 509–542 (2019). Publisher: Taylor & Francis _eprint: 10.1080/05704928.2018.1506799.

28. K. Chin, C. Wieslander, H. Shi, et al., “Pelvic Organ Support in Animals with Partial Loss of Fibulin-5 in the Vaginal Wall,” PLoS ONE 11, e0152793 (2016).

29. C. K. Wieslander, D. D. Rahn, D. D. McIntire, et al., “Quantification of Pelvic Organ Prolapse in Mice: Vaginal Protease Activity Precedes Increased MOPQ Scores in Fibulin 5 Knockout Mice,” Biol. Reproduction 80, 407–414 (2009).

30. J. P. R. Day, G. Rago, K. F. Domke, et al., “Label-free imaging of lipophilic bioactive molecules during lipid digestion by multiplex coherent anti-Stokes Raman scattering microspectroscopy,” J. Am. Chem. Soc. 132, 8433–8439 (2010).

31. Z. Movasaghi, S. Rehman, and I. U. Rehman, “Raman Spectroscopy of Biological Tissues,” Appl. Spectrosc. Rev. 42, 493–541 (2007). Publisher: Taylor & Francis _eprint: 10.1080/05704920701551530.

32. K. A. Dehring, A. R. Smukler, B. J. Roessler, and M. D. Morris, “Correlating changes in collagen secondary structure with aging and defective type II collagen by Raman spectroscopy,” Appl. Spectrosc. 60, 366–372 (2006).

33. L. Salvatore, N. Gallo, M. L. Natali, et al., “Mimicking the Hierarchical Organization of Natural Collagen: Toward the Development of Ideal Scaffolding Material for Tissue Regeneration,” Front. Bioeng. Biotechnol. 9, 644595 (2021).

34. C. N. Grover, R. W. Farndale, S. M. Best, and R. E. Cameron, “The interplay between physical and chemical properties of protein films affects their bioactivity,” J. Biomed. Mater. Res. Part A 100A, 2401–2411 (2012). _eprint: https://onlinelibrary.wiley.com/doi/pdf/10.1002/jbm.a.34187.

35. W.-T. Cheng, M.-T. Liu, H.-N. Liu, and S.-Y. Lin, “Micro-Raman spectroscopy used to identify and grade human skin pilomatrixoma,” Microsc. Res. Tech. 68, 75–79 (2005).

36. C. Gullekson, L. Lucas, K. Hewitt, and L. Kreplak, “Surface-Sensitive Raman Spectroscopy of Collagen I Fibrils,” Biophys. J. 100, 1837–1845 (2011).

37. L. Tong, Z. Hao, C. Wan, and S. Wen, “Detection of depth-depend changes in porcine cartilage after wear test using Raman spectroscopy,” J. Biophotonics 11, e201700217 (2018). _eprint: https://onlinelibrary.wiley.com/doi/pdf/10.1002/jbio.201700217.

38. Z. Chi, X. G. Chen, J. S. W. Holtz, and S. A. Asher, “UV Resonance Raman-Selective Amide Vibrational Enhancement: Quantitative Methodology for Determining Protein Secondary Structure,” Biochemistry 37, 2854–2864 (1998). Publisher: American Chemical Society.

39. V. Renugopalakrishnan, L. A. Carreira, T. W. Collette, et al., “Non-Uniform Triple Helical Structure in Chick Skin Type I Collagen on Thermal Denaturation: Raman Spectroscopic Study,” Zeitschrift für Naturforschung C 53, 383–388 (1998). Publisher: De Gruyter.

40. M. J. Domingo, A. G. Franques, Q. Zhou, and K. S. Miller, “Biaxial contractility of the murine cervix with elastic fiber deficiency,” (SB3C Foundation, Inc., Vail, Colorado, USA, 2023).

41. D. C. LeBert, J. M. Squirrell, J. Rindy, et al., “Matrix metalloproteinase 9 modulates collagen matrices and wound repair,” Dev. (Cambridge, England) 142, 2136–2146 (2015).

42. H. F. Bigg, A. D. Rowan, M. D. Barker, and T. E. Cawston, “Activity of matrix metalloproteinase-9 against native collagen types I and III,” The FEBS J. 274, 1246–1255 (2007). _eprint: https://onlinelibrary.wiley.com/doi/pdf/10.1111/j.1742-4658.2007.05669.x.

43. M. Mahendroo, “Cervical remodeling in term and preterm birth: insights from an animal model,” Reproduction 143, 429–438 (2012). Publisher: BioScientifica Section: Reproduction.

44. R. Institute of Medicine (US) Roundtable on Environmental Health Sciences, D. R. Mattison, S. Wilson, et al., “Preterm Birth and Its Consequences,” in The Role of Environmental Hazards in Premature Birth: Workshop Summary, (National Academies Press (US), 2003).

45. M. L. Akins, K. Luby-Phelps, and M. Mahendroo, “Second harmonic generation imaging as a potential tool for staging pregnancy and predicting preterm birth,” J. Biomed. Opt. 15, 026020 (2010).

46. K. Yoshida, H. Jiang, M. Kim, et al., “Quantitative Evaluation of Collagen Crosslinks and Corresponding Tensile Mechanical Properties in Mouse Cervical Tissue during Normal Pregnancy,” PLoS ONE 9, e112391 (2014).

47. M. Jastrzebska, R. Wrzalik, A. Kocot, et al., “Raman spectroscopic study of glutaraldehyde-stabilized collagen and pericardium tissue,” J. Biomater. Sci. Polym. Ed. 14, 185–197 (2003). Publisher: Taylor & Francis _eprint: 10.1163/156856203321142605.

48. T. A. Shaik, A. Alfonso-García, X. Zhou, et al., “FLIm-Guided Raman Imaging to Study Cross-Linking and Calcification of Bovine Pericardium,” Anal. Chem. 92, 10659–10667 (2020). Publisher: American Chemical Society.

49. J. Trägårdh, T. Pikálek, M. Šerý, et al., “Label-free CARS microscopy through a multimode fiber endoscope,” Opt. Express 27, 30055–30066 (2019). Publisher: Optica Publishing Group.

50. A. Lombardini, V. Mytskaniuk, S. Sivankutty, et al., “High-resolution multimodal flexible coherent Raman endoscope,” Light. Sci. & Appl. 7, 10 (2018). Number: 1 Publisher: Nature Publishing Group.

51. H. Bae, M. Rodewald, T. Meyer-Zedler, et al., “Feasibility studies of multimodal nonlinear endoscopy using multicore fiber bundles for remote scanning from tissue sections to bulk organs,” Sci. Reports 13, 13779 (2023). Number: 1 Publisher: Nature Publishing Group.

52. Y. Zhang, M. L. Akins, K. Murari, et al., “A compact fiber-optic SHG scanning endomicroscope and its application to visualize cervical remodeling during pregnancy,” Proc. National Acad. Sci. 109, 12878–12883 (2012). Publisher: Proceedings of the National Academy of Sciences.

