## Supplemental figures for "Collagen organization and structure in *FLBN5*^-/-^, mice using label-free microscopy: implications for pelvic organ prolapse"

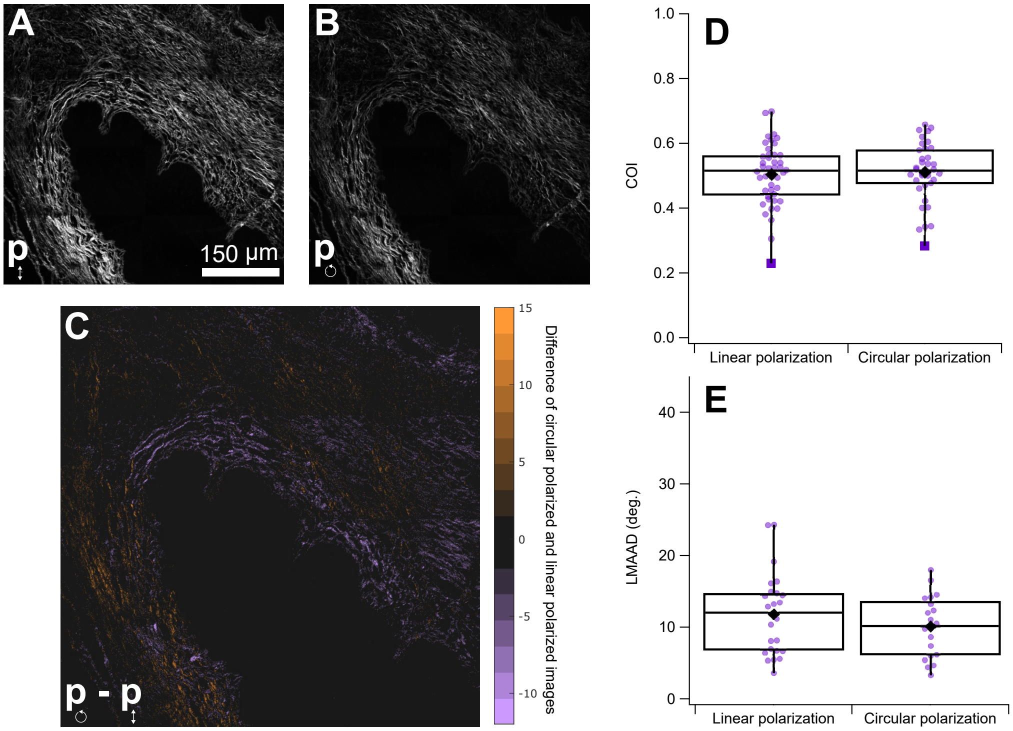


**Fig. S1.** Effect of laser polarization on SHG imaging of POP tissue. SHG images of the same region of interest using a linearly polarized (**A**) and circularly polarized (**B**) excitation beam. Linear polarization induces a greater second harmonic intensity response than circular polarization at the same input laser power. Circularly polarized light was generated by inserting a $\lambda/4$ wave plate (AQWP05M-980, Thorlabs) in the beam path of the linearly polarized beam. After median normalization of each image, a difference image of the circularly polarized and linearly polarized images was calculated (**C**) to highlight the effect of polarization on the second harmonic response of collagen. In the difference image, orange pixels highlight collagen with relatively greater second harmonic response to circularly polarized light than linearly polarized light. Conversely, purple pixels show greater second harmonic response to linearly polarized light. We quantified the effect of linear polarized and circular polarized light on the COI and LMAAD morphology metrics and observed no significant difference (**D** and **E**).

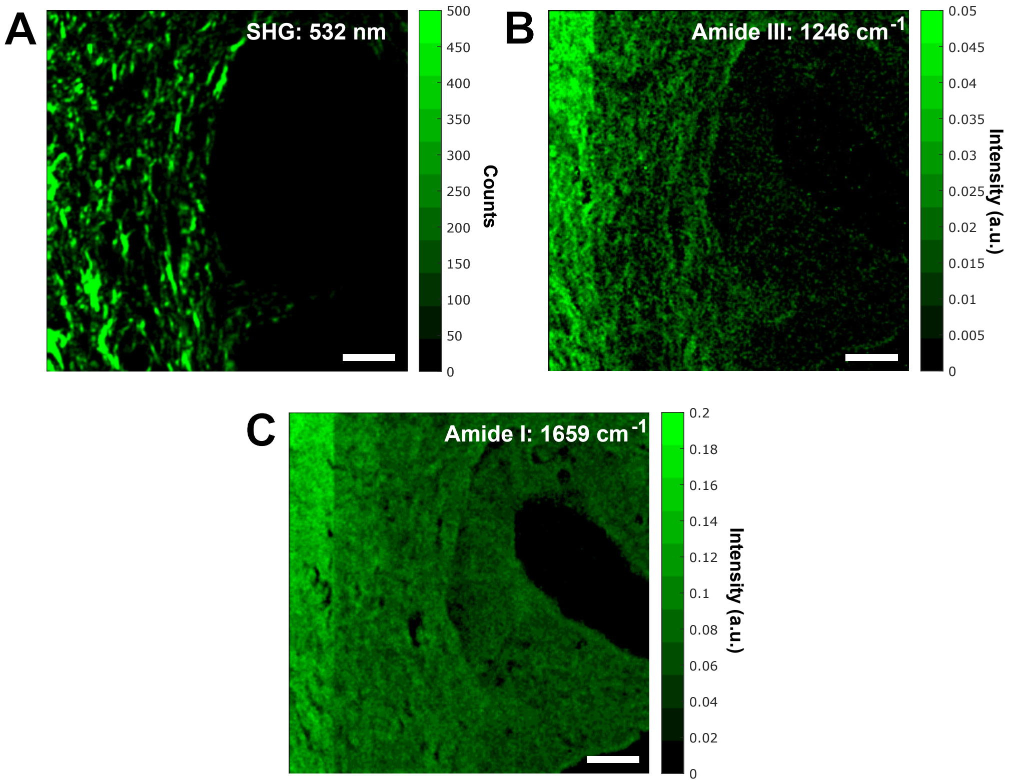


**Fig. S2.** The amide III band contains vibrational modes that are more specific to collagen than the amide I band. (**A**) Using the BCARS pump laser (1064 nm), second harmonic signal (532 nm) from collagen was measured using a spectrometer. On the same region, a BCARS hyperspectral image was acquired using the same pixel spacing as the SHG image. (**B-C**) An intensity projection at 1246 $cm^{-1}$ of the amide III is more similar to the SHG image than an intensity projection at 1659 $cm^{-1}$ of the amide I. The scale bar is 25 µm.

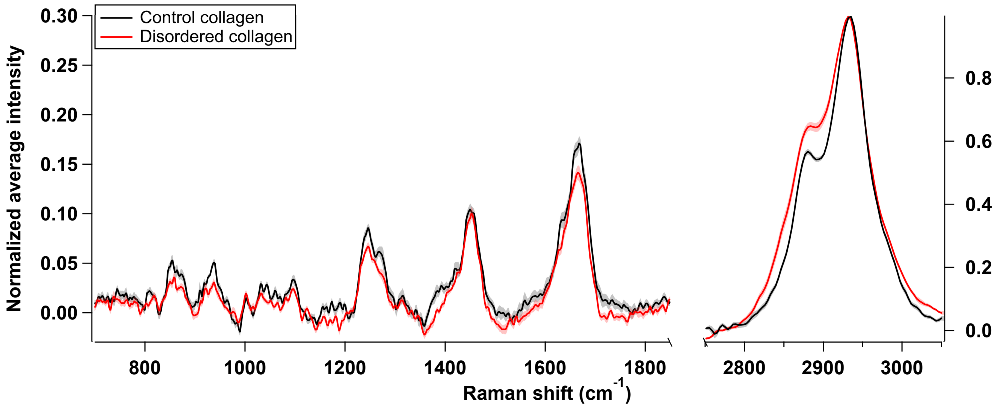


**Fig. S3.** BCARS spectra of temperature-induced changes to the secondary structure of *in-vitro* collagen gels. Following pH neutralization, collagen gels (5 mg/ml, Collagen Type I, Rat Tail, Ibidi) were polymerized for 20 hours at 4°C. Following polymerization, the disordered sample was prepared by placing the gel on a microscopy slide on a hotplate at 65°C for five minutes. The control and disordered samples were imaged via BCARS microscopy in several locations (Control, n = 10, Disordered, n = 14). Error shadows represent the standard error. The amide III region (1220 – 1300 $cm^{-1}$) is a readout of the secondary structure of collagen. The amide III of the control collagen gel contains a right shoulder (at 1271 $cm^{-1}$) that indicates the native triple helix of collagen, which is substantially reduced in the disordered collagen gel.
